# Time-dependent changes to sepsis-specific networks in the plasma proteome are mechanistic readouts of sepsis progression

**DOI:** 10.1101/2020.09.08.285221

**Authors:** G. Pimienta

**Affiliations:** Proteomics Core, NIH grant HL131474, Sanford Burnham Prebys Medical Discovery Institute, La Jolla, CA 9207

**Keywords:** Plasma proteomics, protein network, sepsis

## Abstract

Sepsis accounts for 1 in 5 deaths globally and is the most common cause of deaths in U.S. hospitals. Despite this public health burden, no diagnostic biomarker, nor therapeutic agent for sepsis has proven useful or effective. The principal obstacle is the lack of a mechanistic understanding of this syndrome, particularly during its onset and progression. Using an experimental model of murine sepsis, we report here a time-dependent assessment of changes to the plasma proteome upon infection with *Salmonella enterica* serovar Typhimurium. Changes to the plasma proteome signature of sepsis (PPSS) revealed a transition from early inflammation and coagulation to a later stage of chronic inflammation, coagulopathy and bacteremia. This study represents an advance in our understanding of sepsis progression that may guide innovative therapeutic attitudes and help clinicians track sepsis progression.

## INTRODUCTION

Sepsis is a life-threatening disease due to a dysregulated host response to infection with induction of tissue coagulopathy and inflammation [1]. Sepsis accounts for 1 in 5 deaths globally and is the most common cause of deaths in U.S. hospitals [2]. Among patients with severe sepsis or septic shock, mortality averages 25-30% [3] with many survivors experiencing long-term cognitive impairment, residual organ damage and recurrent infections [4]. Despite this public health burden, no diagnostic biomarker, nor therapeutic agent for sepsis has proven useful or effective [5–7]. This failure to diagnose and treat sepsis is a strong indication that our current mechanistic understanding of this syndromic immune disorder is incomplete, or that the therapy trial approaches have so far been flawed [5–7]. Our current understanding of human sepsis progression rests on the sequential appearance of two clinical stages – early and late sepsis –, defined by clinical symptoms and correlational changes in single protein abundances in blood, largely devoid of a mechanistic explanation [8]. So far, we understand that early sepsis is triggered by a near simultaneous amplification of immune activation and suppression mechanisms, and coagulation activity at the site of infection [9–10]. If the exacerbated immune-hemostatic response remains unresolved, its self-amplifying intensity leads to late sepsis, characterized by disseminated intravascular coagulopathy, multiple organ damage and possible death [9–10]. How and which biochemical mechanisms cooperate to drive the onset of early sepsis and the transition to late sepsis is still an open question.

The cellular and secreted proteome composition at any given time is, together with the metabolome, the direct mediator of the biochemical functions that dictate cellular and organismal phenotype [11]. Most importantly, proteins do not act in isolation, but as networks [12] with a context-dependent protein node stoichiometry [13]. The motivation of this study rests on the hypothesis that measurable changes to sepsis-specific protein networks in the plasma proteome will help track sepsis progression and provide a mechanistic framework of the events that trigger the transition from early to late sepsis. To test this hypothesis, we have assessed abundance changes to the mouse plasma proteome at six time points, upon infection with *Salmonella enterica* serovar Typhimurium (ST), a murine model of sepsis optimized in our laboratory [14], and which we have recently used to integrate the Plasma Protein Signature of Sepsis (PPSS), a network of 84 secreted proteins that have a known role in human sepsis [15]. Our results show a two-step reconfiguration of the PPSS network. The first change occurred on day 4 and was indicative of an exacerbated immune-hemostatic burst in activities that remained unresolved and at high abundances throughout the following time points (days 5-8). The second change to the PPSS appeared on day 5, was concomitant to bacteria in blood, and indicative of consumptive coagulopathy and complement exhaustion. Observations of this report are interpreted in the context of clinical observations on human sepsis.

## MATERIALS AND METHODS

### Mouse infection

We have previously described the infection protocol used in this study [14]. In brief, 12-week old sterile male mice (strain C57BL/6J) were infected via oral gavage with 1 × 10^7^ colony formation units (CFU) in sterile phosphate buffer saline solution of the Gram-negative bacteria Salmonella enterica serovar Typhimurium (ST) reference strain ATCC 14028 (CDC 6516-60). Four uninfected mice were sacrificed as controls, prior to infection (day 0). Four additional mice per day were sacrificed on days 2, 3, 4, 5 and 8 (late sepsis). We collected 200-500 μL of whole blood in analytical tubes coated with EDTA (Fisher Scientific) by cardiopuncture of anesthetized mice. To calculate colony forming units (CFU), a 1:100 dilution of blood in sterile PBS was spread onto ST-specific agar plates, Brilliant green/XLD (Fisher Scientific 2140S86) and incubated overnight at 37 °C. Colonies were counted after 18 h and CFU/mL in blood was calculated. Platelet-poor plasma was prepared in a two-step centrifugation of whole blood at high speed (14,000 rpm), discarding the pellet each time. Plasma samples were mixed with protease inhibitors (Roche) and stored in 20 μL aliquots at −80 °C, until use.

### Proteomics plasma sample preparation

Proteomics samples were prepared as described previously [15], with the exception of the depletion step, which we have optimized as described below. Albumin and Immunoglobulin depletion were performed by precipitating 10 μL of plasma with pre-chilled acetone (20 °C), air-dried and incubated for an hour at 4 °C with a mix of antibodies specific to Albumin (SigmaAldrich LSKMAGL10) and IgA/G (ThermoFisherScientific 88802), crosslinked to magnetic beads. The eluate was collected and incubated once more with magnetic beads. Protein eluates were cysteine-reduced and alkylated with DTT and iodoacetamide, respectively, followed by overnight digestion with mass spectrometry grade Trypsin protease (PIERCETM 90057). Equal amounts of protein digests from the six time points were labeled with specific isotopically labeled tandem mass tags (TMT). Each time point included plasma from two biological replicates (two mice).

### Data collection, analysis and interpretation

The TMT-labeled sample from each biological replicate was injected three times (to generate three technical replicates) in a nano-Liquid chromatography system (nLC) connected inline to a LUMOS-Trybrid Orbitrap tandem mass spectrometer (MS/MS). To increase the number of peptides targeted for sequencing by MS/MS fragmentation, we used a two-dimensional (2D) nLC pre-fractionation setup. The first dimension was a low resolution C18 reverse-phase separation at high pH that produced 12 fractions. Each fraction was in turn separated at low pH in a high resolution C8 reverse phase and ionized for MS/MS fragmentation and MS determination. The separation and data collection methods were as described previously [15]. Data Protein identification assignments and TMT ratio calculations were performed using the latest version of MaxQuant (1.6.8) [16]. Statistical calculations and plots were done in RStudio (1.1.463). To identify and quantify bacterial and host protein abundance simultaneously, we combined the most updated proteome databases from UniProtKB for Salmonella enterica serovar Typhimurium 14028 and the mouse strain C57BL/6J.

## RESULTS

### Experimental approach and data quality

The goal of this study was to quantify time-dependent changes to the PPSS network in the plasma proteome upon ST infection. The experimental workflow comprised an assessment of bacterial colony forming units (CFU) in blood over an eight-day infection time course and the generation of platelet-poor plasma for multiplex quantitative proteomics. Plasma from each sample underwent protein chemical derivatization, trypsin digestion and peptide N-terminus TMT labeling, followed by mass spectrometry data collection and downstream data analysis. Downstream bioinformatics entailed protein identification, relative abundance quantification and the assessment of time-dependent changes to PPSS network stoichiometry in the plasma proteome (Figure 1).

**Figure 1.**
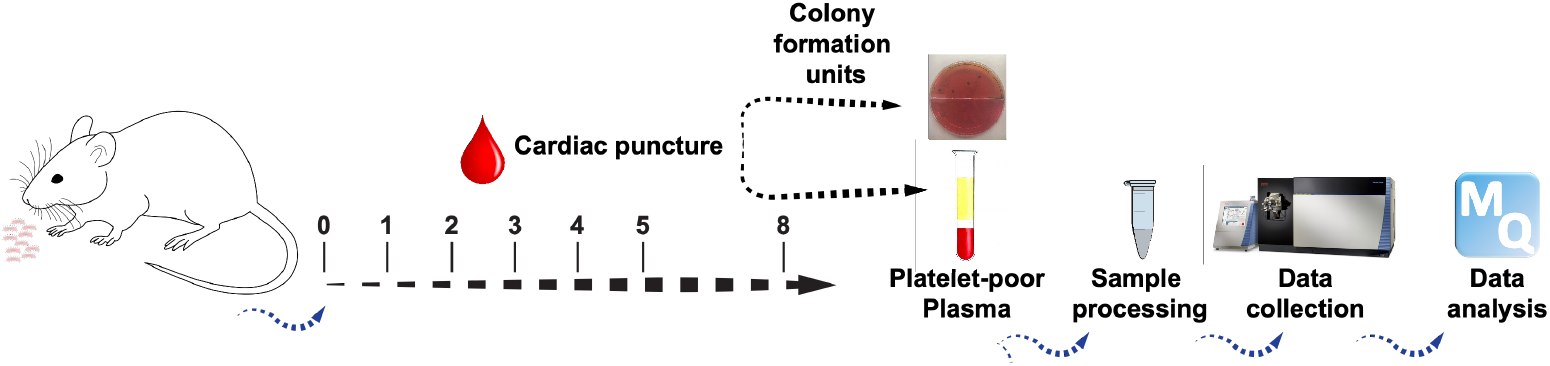
Experimental Design. **A. Experimental design**. Blood was collected by cardiopuncture from 4 mice at each time point and used to prepare platelet-poor plasma for proteomics experiments or to calculate bacterial colony forming units. Plasma proteins from two independent mice at each time point were chemically derivatized, digested with Trypsin, and each “barcoded” with a different TMT label. Once labeled, equal amounts of each time point, was mixed and analyzed by proteomics. Protein identification and relative protein abundances were determined using the proteomics software MaxQuant.

Proteomics data from two mice (biological replicates) was collected at six time points: uninfected mice (control day 0, CTL), and on days 2, 3, 4, 5 and 8, post-infection. The combined dataset from the two biological replicates included fragmentation spectra for 8430 unique peptides, from which 919 proteins were identified with an average identification score of 116 and an average 25% coverage (Supplementary File 1). Technical repeats (three sample injections per biological replicate were highly concordant when compared to each other, with Pearson Correlation Calculation (PCC) values ≥ 0.95 (Supplementary Figs. 1-3). Certainty in protein abundance changes could be assigned within a narrow window: −0.5 ≤ and ≥ 0.5 Day/CTL ratio (Log_2_), based on two-sample T-test p-value calculations (Supplementary Figure 4). When compared, the correlation between the two biological replicates showed that the fold-change values per protein had in most cases the same direction (up or downregulation), as expected for relative quantification in a shotgun experiment (Supplementary Figure 5).

### Changes to the plasma proteome

The plasma proteome dataset comprised ~40% of secreted proteins and ~60% of intracellular proteins, typically regarded as proteinaceous leakage from cell debris [17]. The protein leakage sub-proteome which was composed primarily by intracellular metabolic proteins (Supplementary File 1), showed variable fold-change values throughout the experiment, most of which had an upward trend (Figure 2A). Noticeably, ~80 of these proteins were significantly elevated on days 4-8, including a protein network of 10 proteins retrieved from data curated in STRING as a potential signature of acute liver damage (ALDH1L1, ALDOB, ARG1, ASL, ASS1, BHMT, FAH, FBP1, HPD and MAT1A) (Figures 2A-B) [18–20]. The secreted sub-proteome had two protein dynamics clusters with antiparallel fold-change directionality. The upward cluster comprised ~120 proteins that showed a significant increase in abundances on day 4 that persisted at similar levels throughout the following time points (days 5-8) (Figure 2C and Supplementary Figure 6A). The downward cluster included ~100 secreted proteins that followed a linear downregulation trend throughout days 2-4, before reaching their lowest fold-change values on day 5 (Figure 2D and Supplementary Figure 6B).

**Figure 2.**
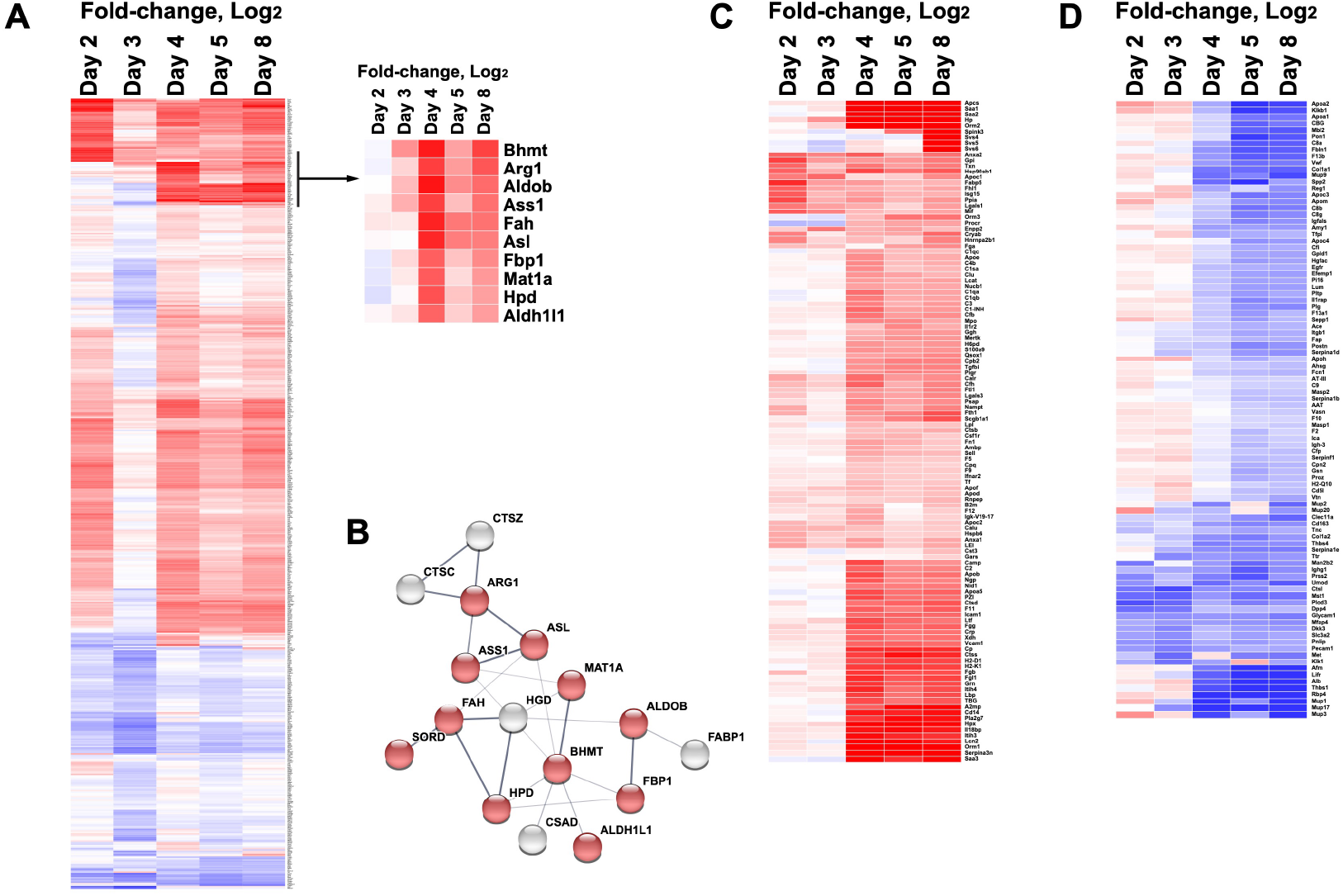
Time-dependent abundance changes to the plasma proteome. **A. Intracellular sub-proteome**. Heatmap representation of the fold-change changes (Log_2_) of the protein leakage sub-proteome, upon unsupervised hierarchical clustering. The heatmap in the inset corresponds to the acute liver damage signature proteins. Heatmap representation the fold-changes (Log_2_) of the putative signature of acute liver damage, upon unsupervised hierarchical clustering. **B**. **Signature network connectivity**. Network extracted by STRING from the proteins with significant changes throughout days 4-8 in the intracellular sub-network (inset in Figure 2B). Protein nodes colored red are those identified as an acute liver signature by STRING, based on publications retrieved from annotation databases [18–20]. Protein nodes colored white interact with the acute liver damage signature but have so far not been associated with liver damage. **C. Upward cluster in the secreted sub-network**. Heatmap representation of the proteins upregulated (fold-change, Log_2_) in the secreted sub-network, upon unsupervised hierarchical clustering. **D. Downward cluster in the secreted sub-network**. Heatmap representation of the proteins downregulated (fold-change, Log_2_) in the secreted sub-network, upon unsupervised hierarchical clustering.

### The PPSS network

From the 84 protein nodes that integrate the PPSS network [15], 14 were not identified in this study (Supplementary Table and Supplementary File). The absence of these proteins in the current dataset was not surprising, given their known low abundance in plasma and in most cases their low molecular weight (Supplementary Table and Supplementary File), two factors known to compromise their reproducible identification in traditional shotgun approaches [21]. Nevertheless, most of the PPSS proteins identified in this study had good identification parameters and a fold-change magnitude on day 8 similar to the one from our previous publication, which was also determined on day 8 in the same mouse model (Figure 3A and Supplementary Table and Supplementary File). The proteins with a good correlation also had a significant reproducibility in the two biological replicates (Figure 3B) and significant p-values (Figures 3C-D).

**Figure 3.**
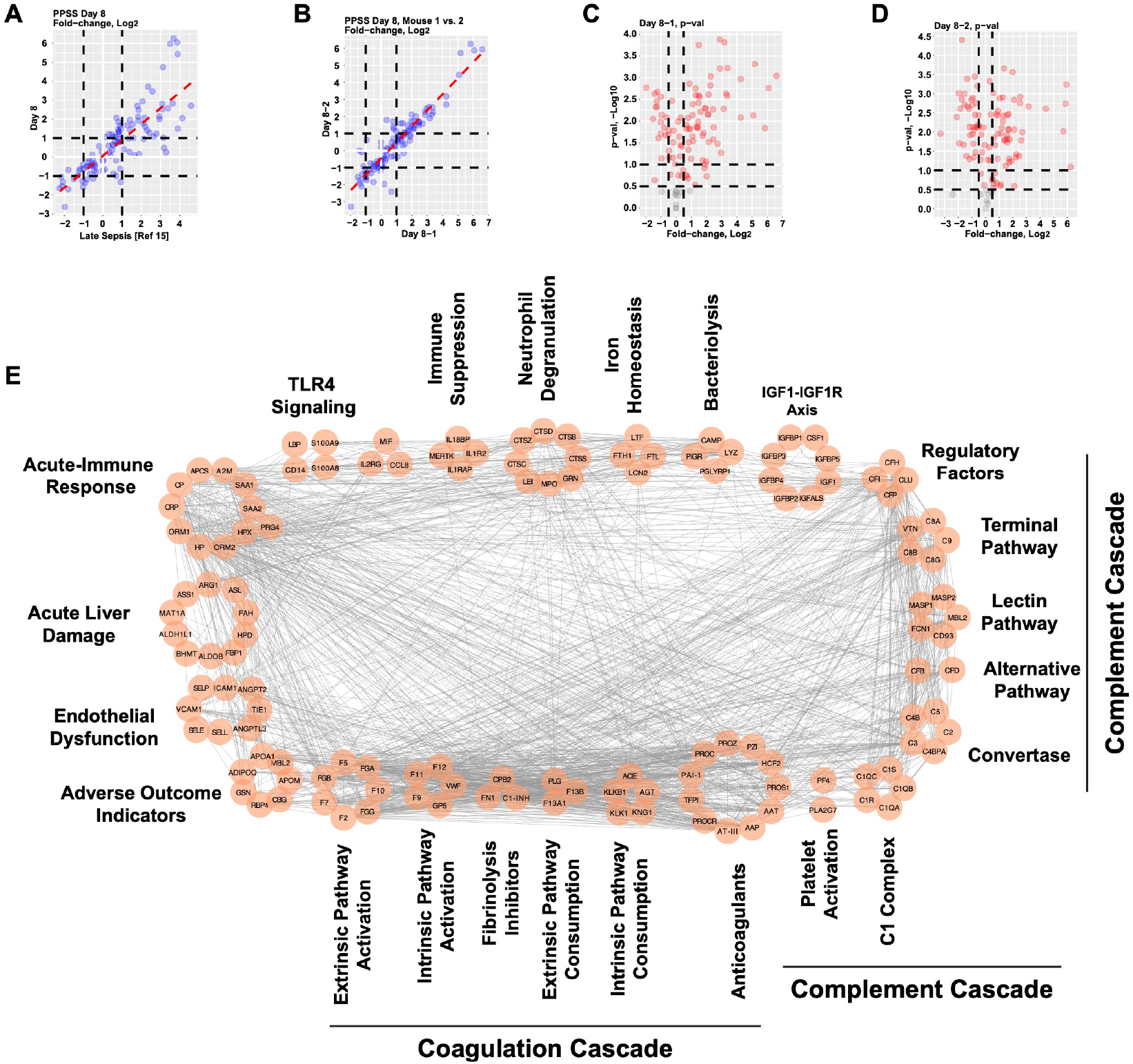
Inspection of PPSS fold-change quality and protein node integration. **A. Fold-change correlation across different experiments**. Scatterplot showing the PPSS fold-changes on day 8 obtained in this study (y axis) when compared to those in our previous publication in late sepsis also on day 8 (x axis). Dashed line colored red is the linear regression, based on the Pearson correlation. **B. Fold-change p-values on day 8**. Scatterplot showing the PPSS fold-changes on day 8 in the two biological replicates obtained in this study. Dashed line colored red is the linear regression, based on the Pearson correlation. **C. Calculated p-values from three technical replicates for the PPSS network in the two biological replicates**. Volcano plots showing the p-values associated to the PPSS fold-changes in the first (left panel) and second (right panel) biological replicates. Proteins with a p-value ≥ 1.0 are colored red, whereas those with a lower value are colored black. **D. Revised PPSS network**. **Integration and subnetwork dissection of the revised PPSS network**. The network was built in Cytoscape from the interaction coordinates extracted in STRING. Functional subnetworks are indicated by a surrounding rectangle and named accordingly. PPSS proteins not identified in this study are colored blue.

We sought to increase the mechanistic information encoded in the PPSS subnetworks by adding protein components not considered in our previous study, through a manual inspection of annotated functions and their functional connectivity to PPSS protein nodes. This approach led to the integration of ~40 proteins not included in our previous publication [15]. The integration of these proteins increased the PPSS network’s connectivity and lead to the rearrangement of the six functional subnetworks proposed previously into seven mechanistic subnetworks and four protein signatures indicative of disease progression (Figure 3E). The seven subnetworks were: *i*. TLR4 signaling [22]; *ii*. Immune Suppression [9, 10]; *iii*. Neutrophil Degranulation [23, 24]; *iv*. Iron Homeostasis [24]; *v*. Bacteriolytic Proteins [25, 26]; *vi*. Coagulation Pathways [27, 28]; and *vii*. Complement Pathways [29, 30]. The four protein signatures were: *i*. Acute Liver Damage [31]; *ii*. Acute-Immune Response [32]; *iii*. Endothelial Dysfunction [33]; and *iv*. Indicators of Adverse Clinical Outcome in Human Sepsis [34]. Other mechanistic subnetworks not identified in this study, but that are components of the PPSS [15], are IGF1-IGF1R axis and Platelet activation (Supplementary References).

### Changes to the PPSS network

Changes to the PPSS network mirrored those of the secreted sub-proteome (Figures 2D-E) and underwent a two-step reconfiguration with opposite trajectories (Figures 4A-B). The first change appeared on day 4 and entailed a switch-like upregulation of ~50 protein activators of innate immune mechanisms and of the coagulation and complement pathways. These abundances remained persistently high throughout days 5-8, with some proteins reaching fold-change levels as high as 100 times than their abundances on day 0 (Figure 4A, Supplementary Table and Supplementary File). The second change to the PPSS appeared on day 5 and was defined by the downregulation of ~30 proteins that reached abundances 2-8 times lower than their baseline levels on day 0. Most of these proteins were coagulation and complement pathway regulators with functions downstream the pathway activators upregulated on day 4 (Figure 4B, Supplementary Table and Supplementary File). Unlike the switch-like increase in protein levels on day 4, the proteins downregulated on day 5 followed a shallow decline in abundances on days 2-4, that preceded the preceded the drop to a lower abundance plateau throughout days 5-8 (Figs. 4B).

**Figure 4.**
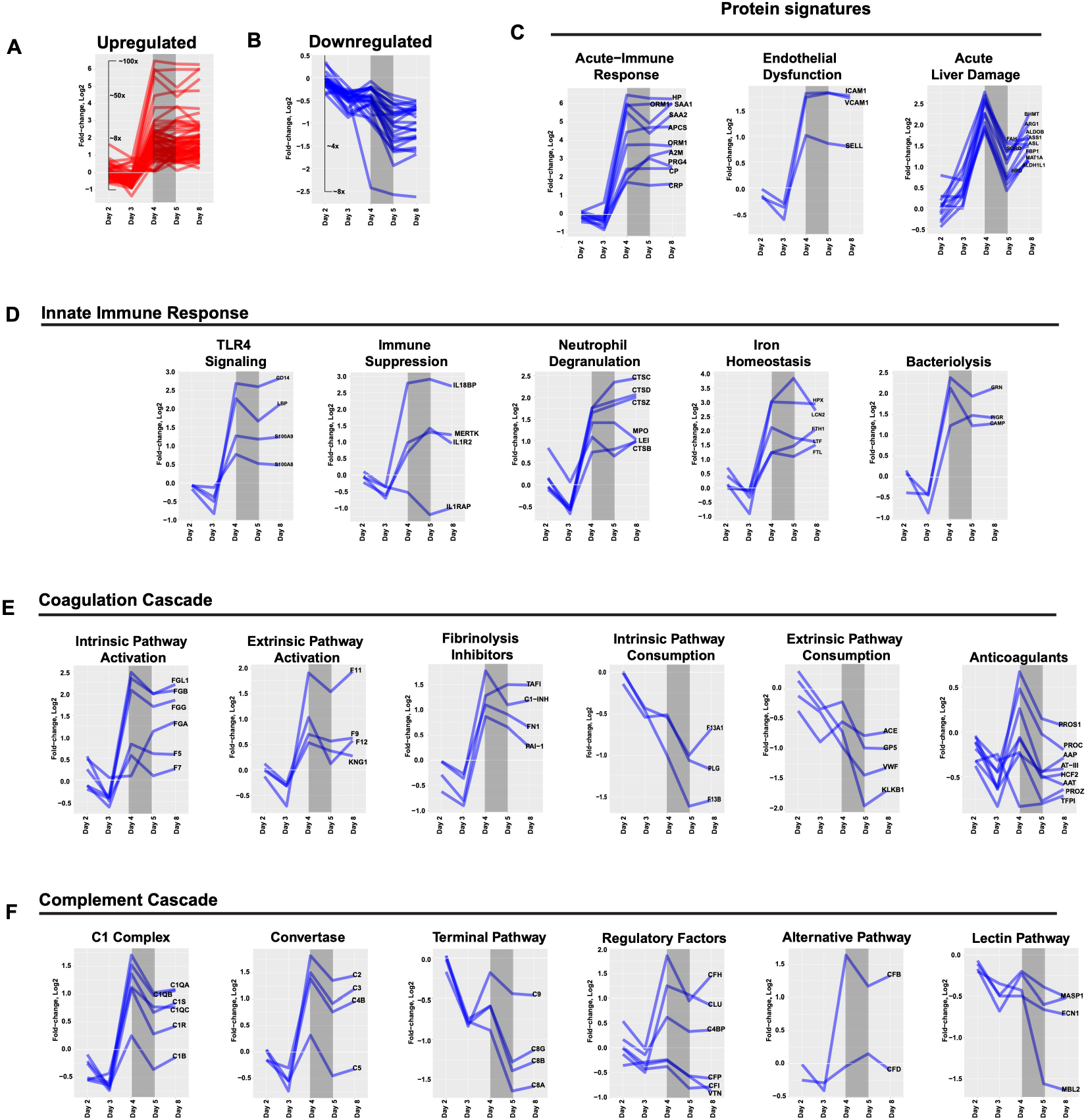
Mechanistic protein dynamics clusters embedded in the PPSS network. **A. Changes to the upward PPSS cluster**. Scatter plot showing the protein dynamics trajectory followed by the PPSS protein nodes upregulated. **B. Changes to the donward PPSS cluster**. Scatter plot showing the protein dynamics trajectory followed by the PPSS protein nodes downregulated. In all panels, the grey inset covering time points day 4 and 5 indicate the transition window from activation of immune-hemostatic processes followed by their burn-out and mechanistic deterioration. **C. Protein signature protein dynamics clusters**. Acute-Immune response kinetic cluster, Endothelial Dysfunction kinetic cluster and Acute Liver Damage kinetic cluster**. D. Innate immune response protein dynamics clusters**. TLR4 signaling, Immune Suppression, Neutrophil Degranulation, Iron Homeostasis and Bacteriolytic Proteins. **E. Coagulation Pathways**. Intrinsic Pathway Activation, Extrinsic Pathway Activation, Fibrinolytic Inhibitors, Intrinsic Pathway Consumption, Extrinsic Pathway Consumption and Anticoagulant Downregulation. **E. Complement Pathways**. C1 complex, C3/C5 Convertase, Terminal Pathway, Complement Pathway Regulators, Lectin Pathway and Alternative Pathway.

Overall, changes to the PPSS network defined a narrow time window on days 4-5 (Figures 4A-B, grey inset) that started with the burst in immune-hemostatic activities on day 4 and was quickly followed on day 5 by the appearance of consumptive coagulopathy and complement exhaustion. Noticeably, groups of protein nodes that participate in the same biochemical process had coordinated time-dependent dynamics and similar fold-change magnitudes, that stood out as subnetwork-specific kinetic clusters (Figures 4C-F). These clusters could be sorted in three functional groups, disease indicators (Figure 4C), innate immune response (Figure 4D), coagulation cascade (Figure 4E) and complement cascade (Figure 4F). The dissection of the PPSS network into kinetic clusters was particularly useful in the deconstruction of the coagulation and complement pathways into the three modular components in these pathways: upstream activation, downstream consumption and signal attenuation (Figures 4E-F). Overall, these kinetic clusters represent a quantitative mechanistic readout indicative of sepsis progression that makes the PPSS a dynamic chart that displayed the onset and progression of sepsis, based on the “lighting up or down” of mechanistic protein signatures.

## DISCUSSION

This study builds up on the Plasma Proteomics Signature of Sepsis (PPSS) integrated from the changes to the plasma proteome at a late stage of sepsis in a murine model previously described by us [15]. We have now added mechanistic information to the PPSS in two ways: *i*. we assessed time-dependent changes to the PPSS at six time points post-infection (days 0, 2, 3, 4, 5 and 8); and *ii*. we increased the PPSS subnetwork connectivity by adding protein nodes not included previously (Figs. 3 and 4). This allowed us to determine a stage of transition from early to late sepsis, based on the abundance dynamics of sepsis-specific mechanistic networks in the plasma proteome and of protein signatures used in the clinic as indicators of disease progression and clinical outcome in human sepsis (Figure 5). Changes to the PPSS network upon infection showed a narrow transition window on days 4-5 that resulted from two reconfiguration stages (Figure 5). The first change to the PPSS was a switch-like upregulation of multiple innate immune and hemostatic activation mechanisms, all of which persisted at similarly high levels beyond day 4 (Figure 4A). The second change to the PPSS appeared on day 5 indicating consumptive coagulopathy and complement exhaustion, coupled to the appearance of bacteria in blood evident mouse sickness (Figure 4B).

**Figure 5.**
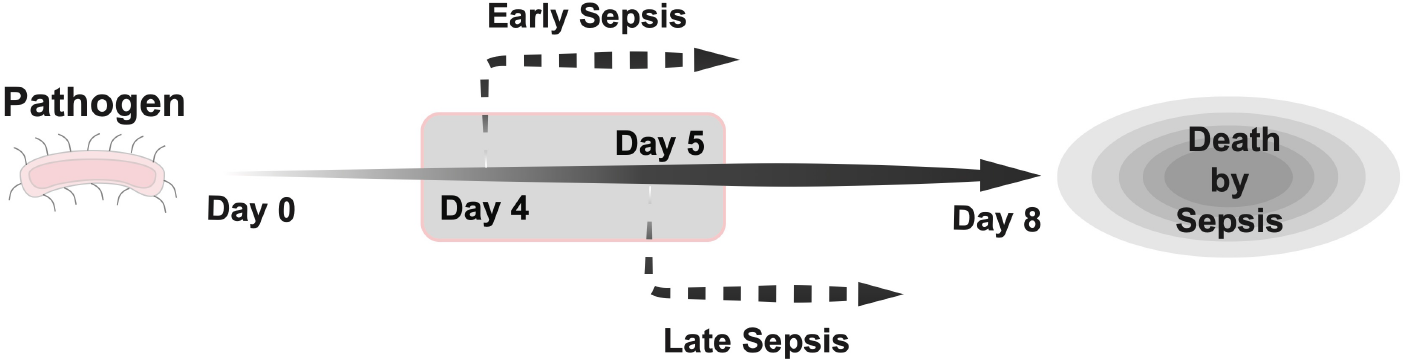
Time-dependent changes to the PPSS network based on a continuous color heatmap. This figure depicts the PPSS network colored in a continuous color gradient that reflects the fold-change magnitudes (Log_2_) specific to each protein node along the six time points assessed. The color gradient is set as blue (downregulated) and red (upregulated) with maximum hue at −3 and 3 (fold-change, Log_2_), respectively. PPSS protein nodes not identified in this study are shown in white and circled in blue.

The significance of this study is two-fold: *i*. the protein nodes embedded in the PPSS all have a documented function in human sepsis (Supplementary Table and Supplementary References) [35]; and *ii*. the protein subnetworks embedded in the PPSS represent mechanistic readouts, each with a unique time-dependent protein abundance trace (Figures 4C-F). This opens the possibility to use the PPSS network or a context-specific version of it, as a time-dependent quantitative landscape to track the onset and progression of experimental sepsis as we have done it here and perhaps also in human sepsis as a replacement or in combination with the symptom-based correlative indicators, currently used in the clinic. We project that the use of the PPSS network in the above-mentioned context will catalyze the development network pharmacology [36, 37], and resolution pharmacology [38, 39] strategies, tailored to halt the short-lived transition from the burst in immune-hemostatic activities to the appearance of coagulopathy and generalized loss of vital functions. To our knowledge approaches of this nature have not been used in the 100-plus clinical trials so far documented. It is therefore possible that a reassessment of these trials by targeting rate-limiting determinants of PPSS subnetworks instead of individual protein mediators, and by planning therapeutic interventions at the right time window, may help reverse impending immune disorder that precedes the irreversible turning point to coagulopathy and organ damage.

## Supporting information

Supplementary References

## SUPPLEMENTARY FIGURES

**Supplementary Figure 1.**
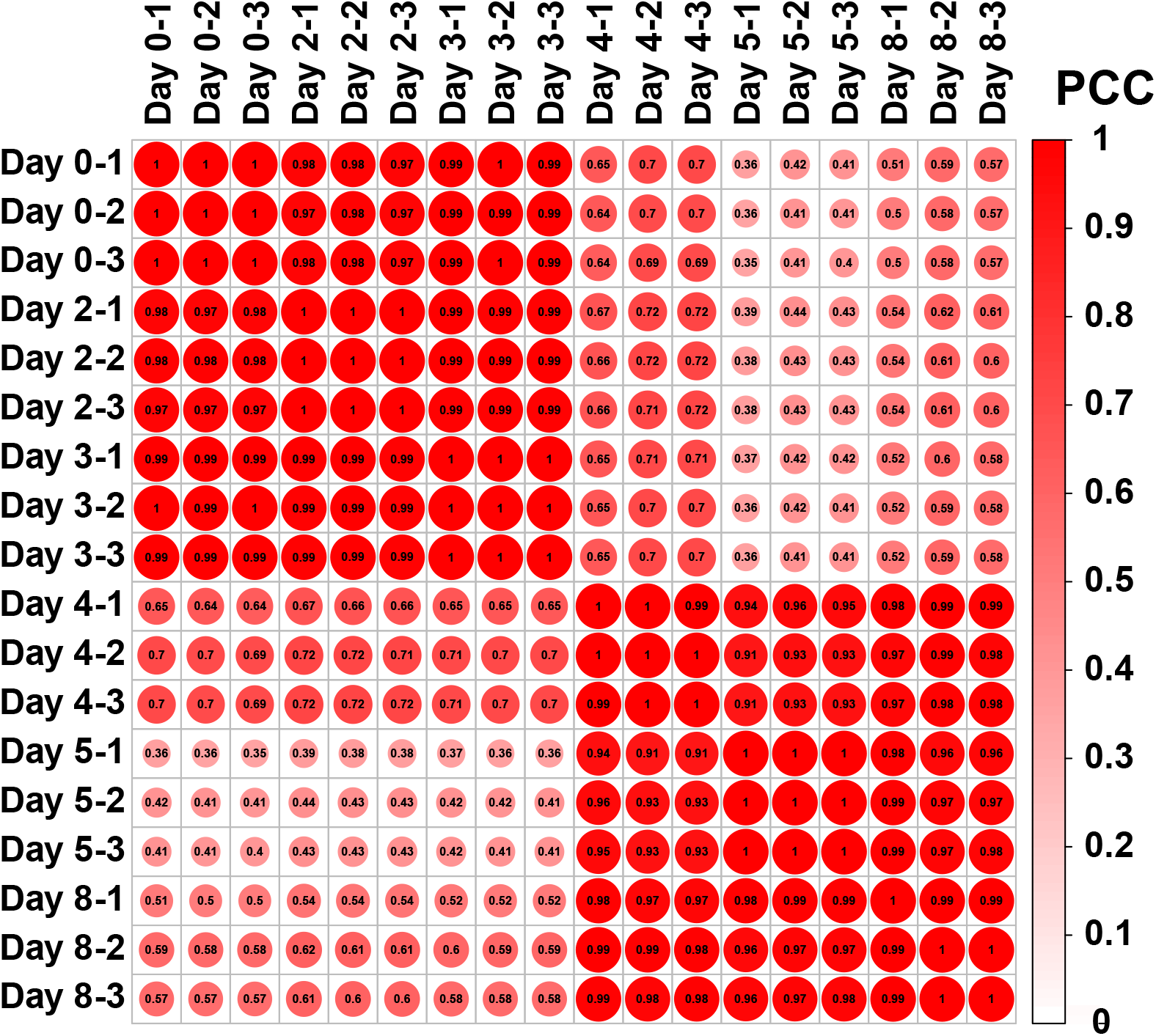
Correlation plot of the Log10 total ion intensity values per protein from each technical replicate per biological replicate. Pearson Correlation Calculation (PCC).

**Supplementary Figure 2.**
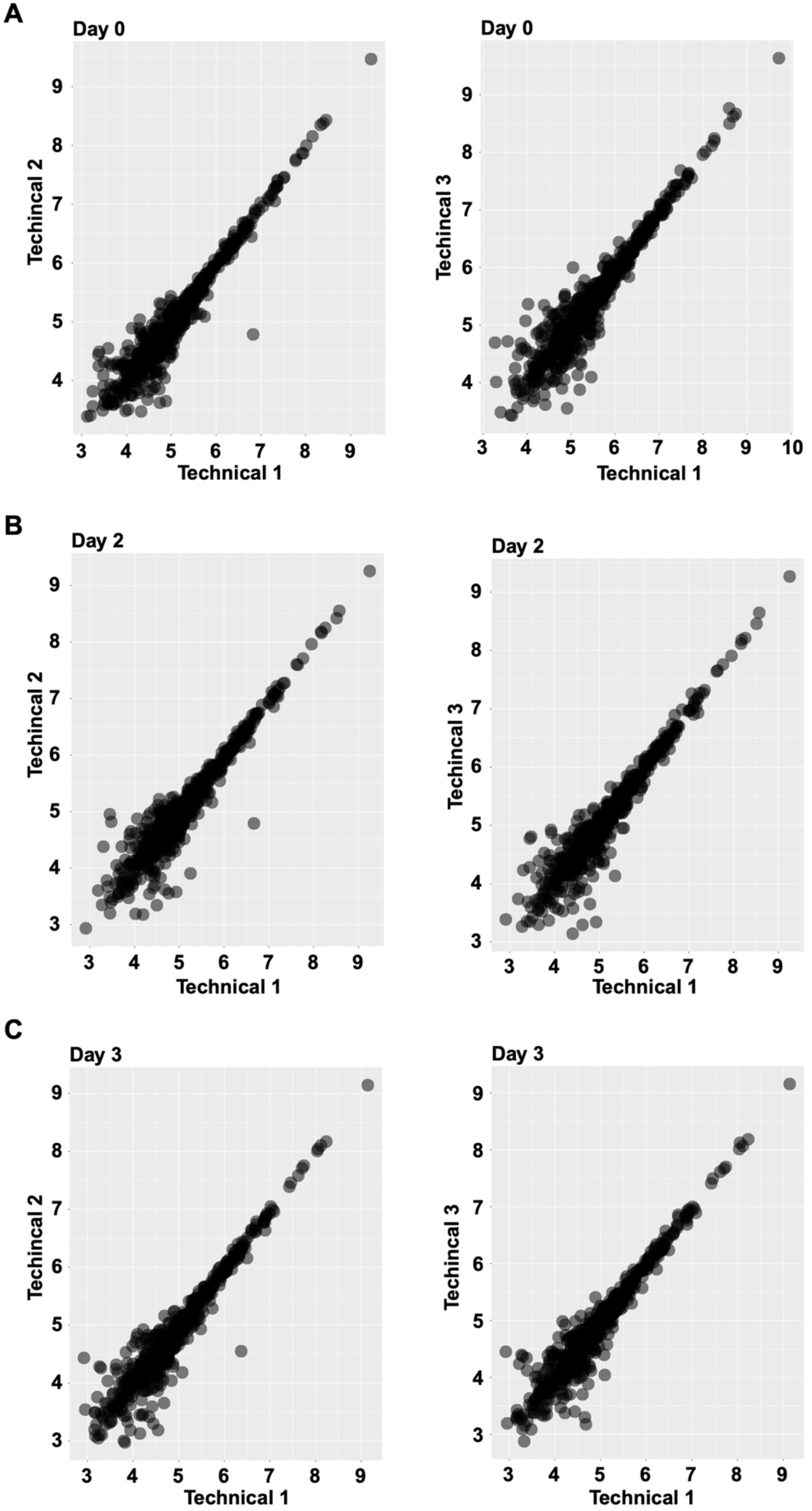
Scatter plots comparing the Log10 total ion intensity values per protein from each technical replicate per biological replicate. Days 0, 2 and 3.

**Supplementary Figure 3.**
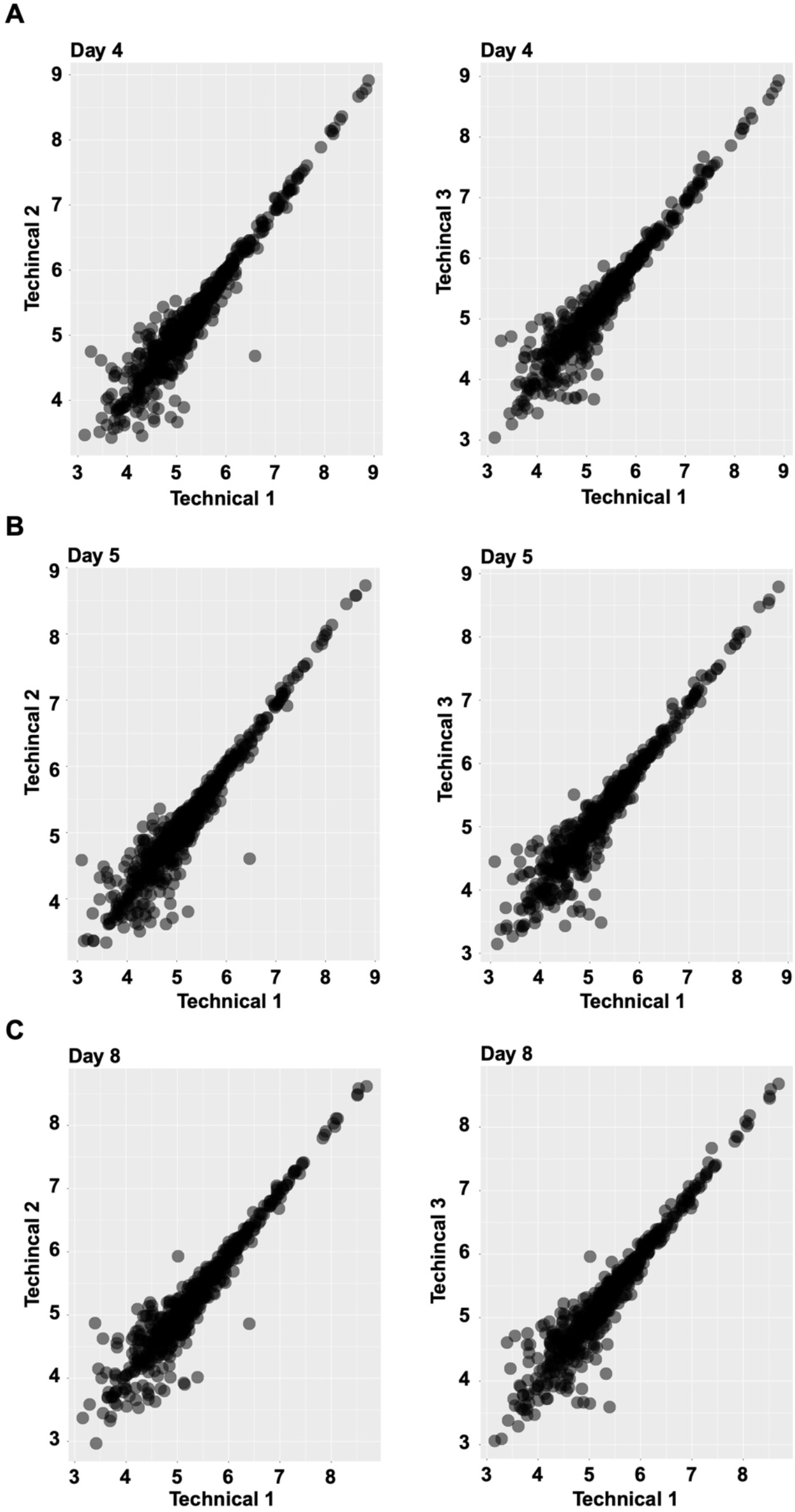
Scatter plots comparing the Log10 total ion intensity values per protein from each technical replicate per biological replicate. Days 4, 5 and 8.

**Supplementary Figure 4.**
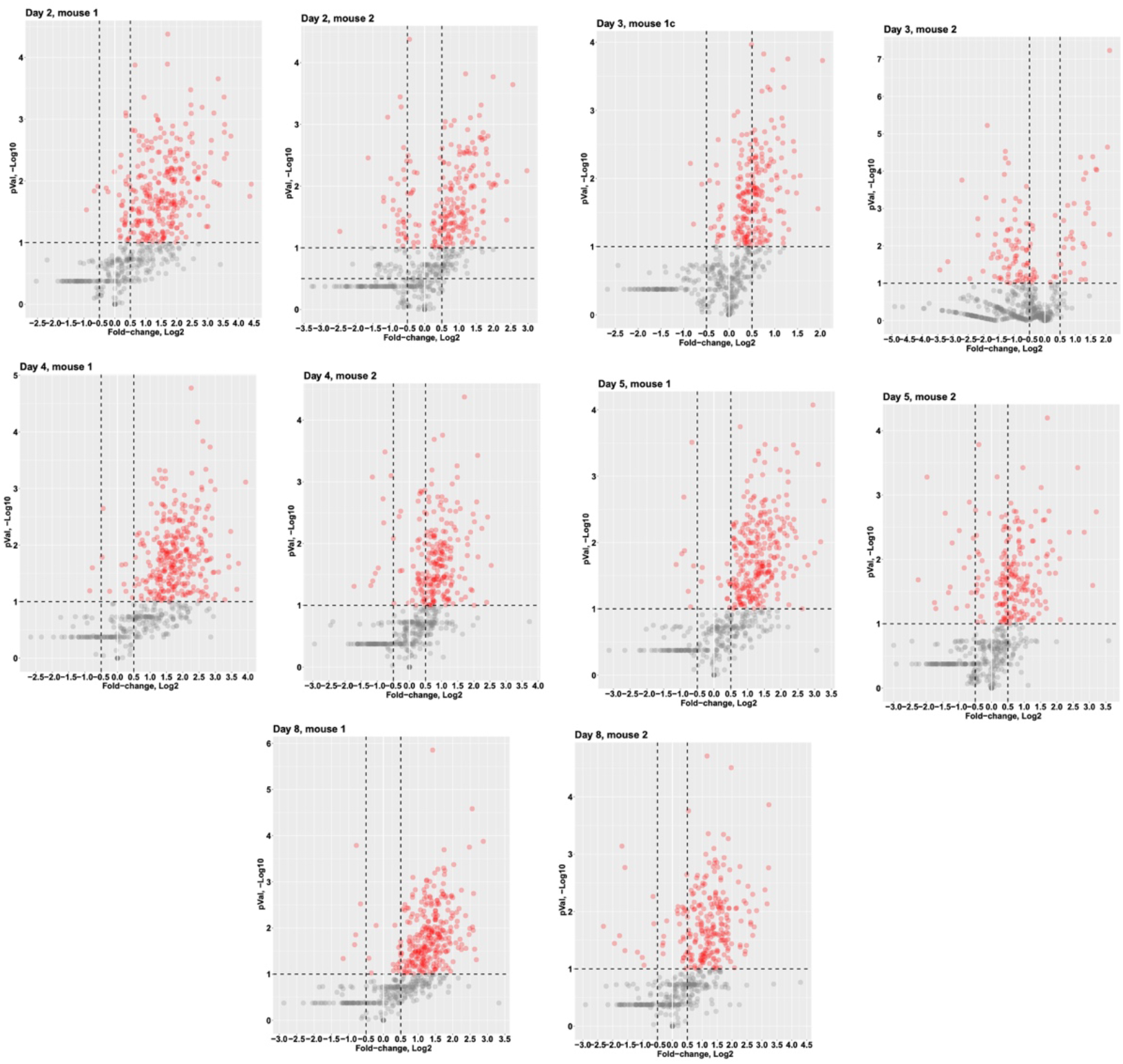
Volcano plots showing −Log10 p-values (y axis) and fold-change values, Log_2_ (x-axis) for each biological replicate. Dots colored red are proteins with a p-value ≥ 1.0, and those grey with values ≤ 1.0.

**Supplementary Figure 5.**
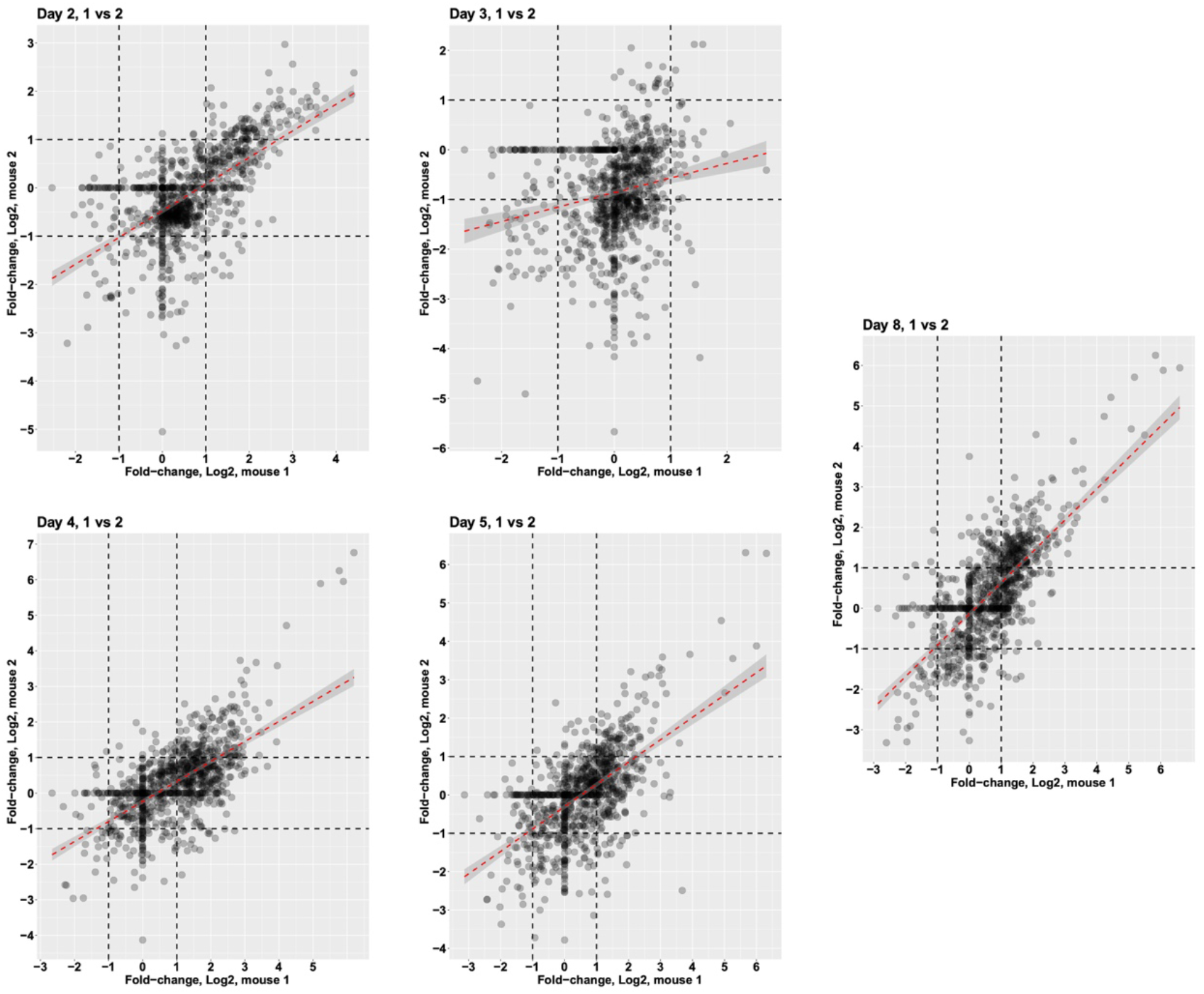
Scatter plots comparing the fold-changes (Log_2_) for each protein identified in the two biological replicates.

**Supplementary Figure 6.**
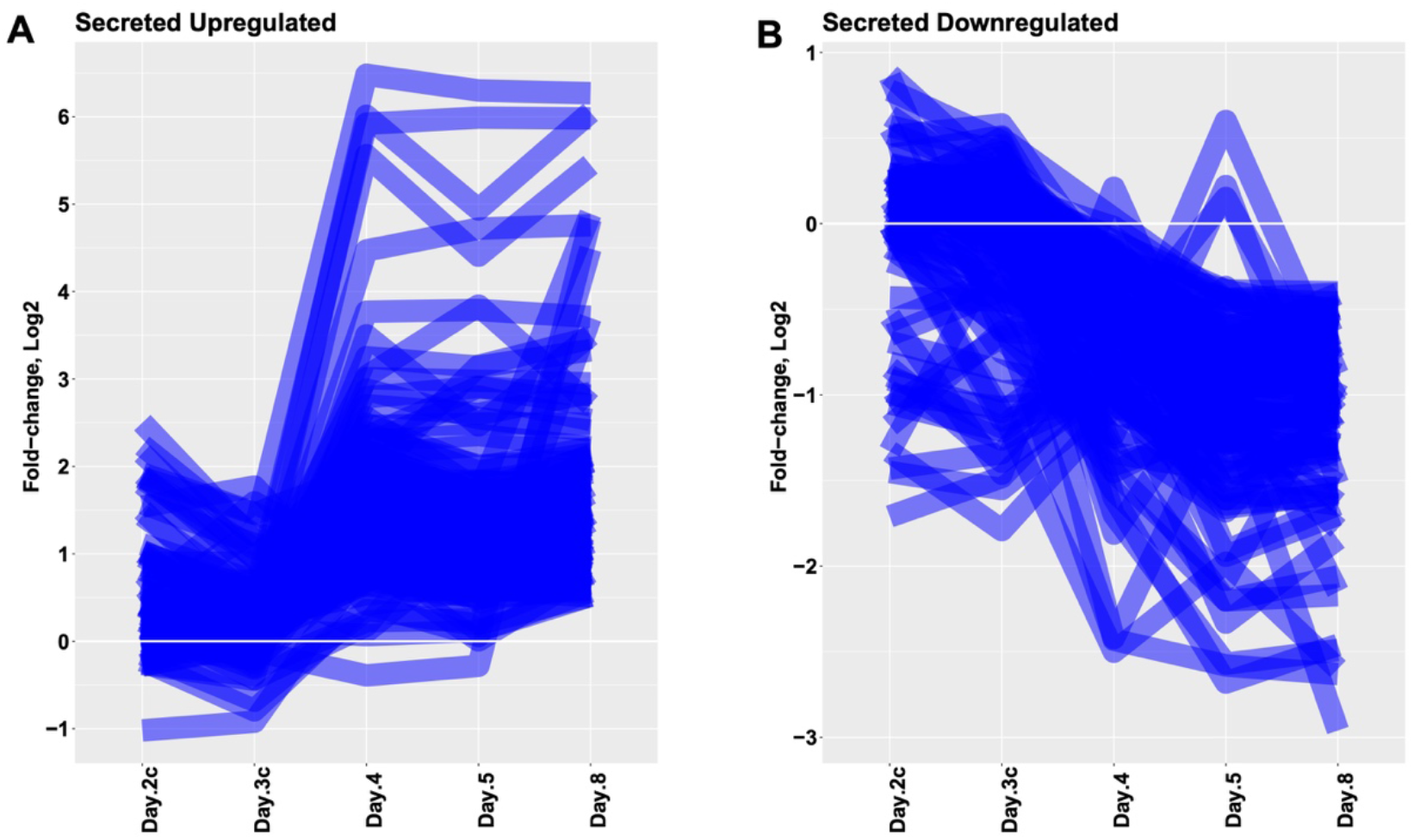
Scatter plots showing the time-dependent fold-changes of the plasma proteins upregulated (panel A) and downregulated (panel B).

## DATASET DEPOSITION

LUMOS-Orbitrap RAW datasets and the results output from the search engine MaxQuant have been deposited in the MassIVE proteomics repository at UCSD and given the identification code PXD021291 at https://massive.ucsd.edu/ProteoSAFe/dataset.jsp?task=1dc72a3bb34a43998f64d20929b61ba3.

## ACKNOWLEDGEMENTS

G.P. is a Staff Scientist working under Jeffrey W. Smith in the Proteomics Core funded by the NIH grant HL131474, at Sanford Burnham Prebys Medical Discovery Institute. The entirety of this report was executed by G.P. when the Proteomics the Core did not have samples to process or data to analyze.

## CONFLICT OF INTEREST

The authors declare no competing financial interest.

## Notes

### Competing Interest Statement

The authors have declared no competing interest.

