## Supplementary References for "Time-dependent changes to sepsis-specific networks in the plasma proteome are mechanistic readouts of sepsis progression"

16. Whalen MJ, Doughty LA, Carlos TM, Wisniewski SR, Kochanek PM, Carcillo JA. Intercellular adhesion molecule-1 and vascular cell adhesion molecule-1 are increased in

the plasma of children with sepsis-induced multiple organ failure. Crit Care Med. 2000;28(7):2600-7. Epub 2000/08/02. doi: 10.1097/00003246-200007000-00070. PubMed PMID: 10921602.

17. Amalakuhan B, Habib SA, Mangat M, Reyes LF, Rodriguez AH, Hinojosa CA, et al. Endothelial adhesion molecules and multiple organ failure in patients with severe sepsis. Cytokine. 2016;88:267-73. Epub 2016/10/05. doi: 10.1016/j.cyto.2016.08.028. PubMed PMID: 27701021; PubMed Central PMCID: PMCPMC5121929.

18. Behnes M, Bertsch T, Lepiorz D, Lang S, Trinkmann F, Brueckmann M, et al. Diagnostic and prognostic utility of soluble CD 14 subtype (presepsin) for severe sepsis and septic shock during the first week of intensive care treatment. Crit Care. 2014;18(5):507. Epub 2014/09/06. doi: 10.1186/s13054-014-0507-z. PubMed PMID: 25190134; PubMed Central PMCID: PMCPMC4174283.

19. Masson S, Caironi P, Fanizza C, Thomae R, Bernasconi R, Noto A, et al. Circulating presepsin (soluble CD14 subtype) as a marker of host response in patients with severe sepsis or septic shock: data from the multicenter, randomized ALBIOS trial. Intensive Care Med. 2015;41(1):12-20. Epub 2014/10/17. doi: 10.1007/s00134-014-3514-2. PubMed PMID: 25319385.

20. Bryant CE, Spring DR, Gangloff M, Gay NJ. The molecular basis of the host response to lipopolysaccharide. Nat Rev Microbiol. 2010;8(1):8-14. Epub 2009/12/01. doi: 10.1038/nrmicro2266. PubMed PMID: 19946286.

21. Dubois C, Marce D, Faivre V, Lukaszewicz AC, Junot C, Fenaille F, et al. High plasma level of S100A8/S100A9 and S100A12 at admission indicates a higher risk of

death in septic shock patients. Sci Rep. 2019;9(1):15660. Epub 2019/11/02. doi: 10.1038/s41598-019-52184-8. PubMed PMID: 31666644; PubMed Central PMCID: PMCPMC6821805.

43. Martensson J, Bell M, Xu S, Bottai M, Ravn B, Venge P, et al. Association of plasma neutrophil gelatinase-associated lipocalin (NGAL) with sepsis and acute kidney

dysfunction. Biomarkers. 2013;18(4):349-56. Epub 2013/05/01. doi: 10.3109/1354750X.2013.787460. PubMed PMID: 23627612.

49. Cenik B, Sephton CF, Kutluk Cenik B, Herz J, Yu G. Progranulin: a proteolytically processed protein at the crossroads of inflammation and neurodegeneration. J Biol

Chem. 2012;287(39):32298-306. Epub 2012/08/04. doi: 10.1074/jbc.R112.399170. PubMed PMID: 22859297; PubMed Central PMCID: PMCPMC3463300.

2010/08/20. doi: 10.1038/ni.1923. PubMed PMID: 20720586; PubMed Central PMCID: PMCPMC2924908.

72

doi: 10.1164/rccm.200709-1400OC. PubMed PMID: 18436791; PubMed Central PMCID: PMCPMC2453509.
