## Supplementary Table for "Time-dependent changes to sepsis-specific networks in the plasma proteome are mechanistic readouts of sepsis progression"

**Fold-change (Day/CTL, $Log_2$). Arrows indicate fold-change magnitude on Day 8. ↑ ≥ 0.5-1.0 ≤, ↑↑ ≥ 1.0-2.0 ≤, ↑↑↑ ≥ 2.0, ↓ -(≥ 0.5-1.0 ≤), ↓↓ -(≥ 1.0-2.0 ≤), ↓↓↓ -(≥ 2.0)**

| **Acute-Phase Responders** | | | | | | | | | | | | | |
|---|---|---|---|---|---|---|---|---|---|---|---|---|---|
| **Fold-change** | **Protein names** | **References** | **Mouse Symbol** | **Human Symbol** | **Day 2** | **Day 3** | **Day 4** | **Day 5** | **Day 8** | **Score** | **Intensity** | **Peptides** | **Mol. weight [kDa]** |
| ↑↑↑ | Alpha-2-macroglobulin-P | | A2mp | A2M | -0.12 | -0.74 | 1.87 | 3.15 | 3.47 | 323.31 | 2.65E+10 | 64.00 | 164.35 |
| ↑↑↑ | Serum amyloid P-component | | Apcs | APCS | 0.06 | -0.30 | 4.46 | 4.72 | 4.75 | 323.31 | 6.41E+10 | 13.00 | 26.25 |
| ↑↑↑ | Ceruloplasmin | | Cp | CP | -0.08 | -0.40 | 2.46 | 2.49 | 2.49 | 323.31 | 4.89E+11 | 71.00 | 121.15 |
| ↑↑ | C-reactive protein | | Crp | CRP | 0.15 | -0.19 | 1.74 | 1.57 | 1.65 | 323.31 | 7.14E+09 | 11.00 | 25.36 |
| ↑↑↑ | Haptoglobin | | Hp | HP | 0.14 | 0.64 | 6.48 | 6.30 | 6.27 | 323.31 | 1.53E+12 | 33.00 | 38.75 |
| ↑↑↑ | Alpha-1-acid glycoprotein 1 | | Orm1 | ORM1 | 0.06 | -0.45 | 3.77 | 3.79 | 3.71 | 323.31 | 3.82E+11 | 18.00 | 23.90 |
| ↑↑↑ | Alpha-1-acid glycoprotein 2 | | Orm2 | ORM2 | -0.23 | -0.11 | 5.92 | 5.99 | 5.98 | 266.75 | 9.09E+10 | 14.00 | 23.84 |
| ↑↑↑ | Serum amyloid A-1 protein | | Saa1 | SAA1 | -0.33 | -0.28 | 6.01 | 4.94 | 6.05 | 323.31 | 1.55E+11 | 10.00 | 13.77 |
| ↑↑↑ | Serum amyloid A-2 protein | | Saa2 | SAA2 | -0.43 | -0.55 | 5.55 | 4.40 | 5.44 | 323.31 | 5.29E+10 | 9.00 | 13.62 |
| ↑↑↑ | Proteoglycan 4 | | Prg4 | PRG4 | -0.20 | -0.86 | 2.29 | 3.03 | 2.58 | 307.96 | 1.85E+09 | 15.00 | 115.99 |
| **Endothelial Dysfunction** | | | | | | | | | | | | | |
| **Fold-change** | **Protein names** | | **Mouse Symbol** | **Human Symbol** | **Day 2** | **Day 3** | **Day 4** | **Day 5** | **Day 8** | **Score** | **Intensity** | **Peptides** | **Mol. weight [kDa]** |
| No | Angiopoietin-2 | | Angpt2 | ANGPT2 | -0.17 | -0.77 | 0.7 | 0.035 | 0.27 | 12.23 | 4.02E+08 | 2.00 | 56.58 |
| ↑↑ | Intercellular adhesion molecule 1 | | Icam1 | ICAM1 | -0.15 | -0.62 | 1.745 | 1.825 | 1.775 | 267.82 | 1.07E+09 | 15.00 | 58.84 |
| ↑ | L-selectin | | Sell | SELL | -0.19 | -0.375 | 1.01 | 0.855 | 0.795 | 48.26 | 1.70E+08 | 7.00 | 42.29 |
| ↑↑ | Vascular cell adhesion protein 1 | | Vcam1 | VCAM1 | -0.025 | -0.3 | 1.82 | 1.835 | 1.725 | 323.31 | 1.98E+09 | 22.00 | 81.32 |
| **Acute liver damage** | | | | | | | | | | | | | |
| **Fold-change** | **Protein names** | | **Mouse Symbol** | **Human Symbol** | **Day 2** | **Day 3** | **Day 4** | **Day 5** | **Day 8** | **Score** | **Intensity** | **Peptides** | **Mol. weight [kDa]** |
| ↑↑ | Cytosolic 10 formyltetrahydrofolate dehydrogenase | | Aldh1l1 | ALDH1L1 | 0.09 | 0.32 | 1.82 | 0.56 | 1.13 | 323.31 | 9.89E+08 | 26.00 | 98.71 |
| ↑↑ | Fructose-bisphosphate aldolase B | | Aldob | ALDOB | 0.00 | 0.80 | 2.76 | 1.35 | 1.74 | 323.31 | 2.56E+09 | 16.00 | 39.51 |
| ↑↑↑ | Arginase-1 | | Arg1 | ARG1 | -0.16 | 0.49 | 2.26 | 1.16 | 1.99 | 188.35 | 1.22E+09 | 14.00 | 34.81 |
| ↑↑ | Argininosuccinate lyase | | Asl | ASL | 0.07 | -0.01 | 2.55 | 1.52 | 1.51 | 155.21 | 8.45E+08 | 16.00 | 51.74 |
| ↑↑ | Argininosuccinate synthase | | Ass1 | ASS1 | 0.18 | 0.94 | 2.24 | 0.76 | 1.67 | 232.92 | 1.27E+09 | 19.00 | 46.58 |
| ↑↑↑ | Betaine--homocysteine S-methyltransferase 1 | | Bhmt | BHMT | -0.11 | 1.29 | 2.68 | 1.06 | 2.22 | 323.31 | 4.23E+09 | 15.00 | 45.02 |
| ↑↑ | Fumarylacetoacetase | | Fah | FAH | 0.28 | 0.26 | 2.59 | 1.57 | 1.65 | 323.31 | 1.00E+09 | 12.00 | 46.18 |
| ↑↑ | Fructose-1,6-bisphosphatase 1 | | Fbp1 | FBP1 | -0.31 | 0.45 | 1.88 | 0.70 | 1.37 | 235.85 | 7.13E+08 | 15.00 | 36.91 |
| ↑↑ | 4-hydroxyphenylpyruvate dioxygenase | | Hpd | HPD | -0.46 | 0.07 | 2.04 | 0.67 | 1.11 | 149.96 | 2.28E+08 | 8.00 | 45.05 |
| ↑↑ | S-adenosylmethionine synthase isoform type-1 | | Mat1a | MAT1A | -0.36 | 0.27 | 1.93 | 0.41 | 1.20 | 311.61 | 3.00E+08 | 6.00 | 43.51 |
| ↑↑ | Sorbitol dehydrogenase | | Sord | SORD | 0.78 | 0.66 | 2.51 | 1.34 | 1.47 | 109.22 | 1.23E+09 | 14.00 | 38.25 |
| **TLR4-signaling** | | | | | | | | | | | | | |
| **Fold-change** | **Protein names** | | **Mouse Symbol** | **Human Symbol** | **Day 2** | **Day 3** | **Day 4** | **Day 5** | **Day 8** | **Score** | **Intensity** | **Peptides** | **Mol. weight [kDa]** |
| ↑↑↑ | Monocyte differentiation antigen CD14 | | Cd14 | CD14 | -0.16 | -0.83 | 2.69 | 2.60 | 2.83 | 323.31 | 4.30E+09 | 14.00 | 39.20 |
| No | Cytokine receptor common subunit gamma | | Il2rg | IL2RG | -1.06 | -0.76 | 0.19 | 0.38 | 0.09 | 11.37 | 8.04E+06 | 2.00 | 42.24 |
| ↑↑↑ | Lipopolysaccharide-binding protein | | Lbp | LBP | -0.07 | -0.50 | 2.28 | 1.67 | 2.13 | 113.25 | 5.71E+08 | 8.00 | 53.06 |
| ↑↑ | Macrophage migration inhibitory factor | | Mif | MIF | 1.64 | 0.85 | 0.82 | 0.65 | 1.06 | 67.33 | 6.17E+08 | 4 | 12.504 |
| No | Protein S100-A8 | | S100a8 | S100A8 | -0.07 | -0.12 | 0.77 | 0.52 | 0.48 | 20.58 | 5.36E+07 | 2.00 | 10.29 |
| ↑↑ | Protein S100-A9 | | S100a9 | S100A9 | -0.05 | -0.38 | 1.27 | 1.18 | 1.24 | 35.56 | 3.40E+08 | 5.00 | 13.05 |
| **Immune suppression** | | | | | | | | | | | | | |
| **Fold-change** | **Protein names** | | **Mouse Symbol** | **Human Symbol** | **Day 2** | **Day 3** | **Day 4** | **Day 5** | **Day 8** | **Score** | **Intensity** | **Peptides** | **Mol. weight [kDa]** |
| ↑↑↑ | Interleukin-18-binding protein | | Il18bp | IL18BP | 0.10 | -0.35 | 2.79 | 2.91 | 2.70 | 197.73 | 6.37E+08 | 3.00 | 21.26 |
| ↑↑ | Interleukin-1 receptor type 2 | | Il1r2 | IL1R2 | -0.22 | -0.60 | 0.68 | 1.42 | 0.96 | 77.77 | 5.19E+08 | 7.00 | 45.65 |
| ↓↓ | Interleukin-1 receptor accessory protein | | Il1rap | IL1RAP | -0.06 | -0.35 | -0.52 | -1.20 | -0.99 | 323.31 | 9.08E+09 | 18.00 | 65.74 |
| ↑↑ | Tyrosine-protein kinase Mer | | Mertk | MERTK | -0.04 | -0.70 | 0.98 | 1.30 | 1.22 | 71.34 | 2.37E+08 | 8.00 | 110.16 |
| **Neutrophil degranulation** | | | | | | | | | | | | | |
| **Fold-change** | **Protein names** | | **Mouse Symbol** | **Human Symbol** | **Day 2** | **Day 3** | **Day 4** | **Day 5** | **Day 8** | **Score** | **Intensity** | **Peptides** | **Mol. weight [kDa]** |
| ↑↑ | Myeloperoxidase | | Mpo | MPO | -0.05 | -0.51 | 1.42 | 1.42 | 1.03 | 38.10 | 7.02E+07 | 6.00 | 81.18 |
| ↑↑ | Cathepsin B | | Ctsb | CTSB | -0.11 | -0.58 | 0.74 | 0.81 | 0.98 | 323.31 | 2.10E+09 | 13.00 | 37.28 |
| ↑↑↑ | Dipeptidyl peptidase 1 | | Ctsc | CTSC | -0.05 | -0.58 | 1.76 | 2.35 | 2.44 | 147.45 | 2.45E+08 | 7.00 | 52.38 |
| ↑↑↑ | Cathepsin D | | Ctsd | CTSD | 0.16 | -0.67 | 1.65 | 1.83 | 2.01 | 137.83 | 6.37E+08 | 9.00 | 44.95 |

| | | | | | | | | | | | | |
|---|---|---|---|---|---|---|---|---|---|---|---|---|
| ↑↑↑ | Cathepsin Z | Ctsz | CTSZ | 0.14 | -0.53 | 1.76 | 1.92 | 2.07 | 79.37 | 4.78E+08 | 7.00 | 34.00 |
| ↑↑↑ | Neutrophil gelatinase-associated lipocalin | Lcn2 | LCN2 | 0.13 | -0.91 | 3.03 | 3.84 | 2.71 | 323.31 | 1.26E+10 | 9.00 | 22.88 |
| ↑↑ | Leukocyte elastase inhibitor A | LEI | SERPINB1 | 0.84 | 0.06 | 1.09 | 0.65 | 1.01 | 61.39 | 2.05E+08 | 9.00 | 42.57 |
| **Iron Homeostasis** | | | | | | | | | | | | |
| **Fold-change** | **Protein names** | **Mouse Symbol** | **Human Symbol** | **Day 2** | **Day 3** | **Day 4** | **Day 5** | **Day 8** | **Score** | **Intensity** | **Peptides** | **Mol. weight [kDa]** |
| ↑↑↑ | Hemopexin | Hpx | HPX | 0.05 | -0.11 | 3.01 | 2.98 | 2.94 | 323.31 | 1.86E+12 | 48.00 | 51.32 |
| ↑↑ | Lactotransferrin | Ltf | LTF | -0.02 | -0.06 | 2.11 | 1.75 | 1.61 | 152.50 | 3.20E+08 | 16.00 | 77.84 |
| ↑↑↑ | Ferritin heavy chain | Fth1 | FTH1 | 0.70 | -0.34 | 1.23 | 1.46 | 2.05 | 66.32 | 3.33E+08 | 3.00 | 21.07 |
| ↑↑ | Ferritin light chain 1 | Ftl1 | FTL | 0.41 | -0.18 | 1.23 | 1.09 | 1.50 | 244.16 | 1.35E+09 | 7.00 | 20.80 |
| **Bactericidal peptides/proteins** | | | | | | | | | | | | |
| **Fold-change** | **Protein names** | **Mouse Symbol** | **Human Symbol** | **Day 2** | **Day 3** | **Day 4** | **Day 5** | **Day 8** | **Score** | **Intensity** | **Peptides** | **Mol. weight [kDa]** |
| ↑↑ | Cathelin-related antimicrobial peptide | Camp | CAMP | -0.38 | -0.41 | 2.14 | 1.23 | 1.28 | 323.31 | 3.92E+08 | 3.00 | 19.45 |
| ↑↑↑ | Granulins | Grn | GRN | 0.08 | -0.44 | 2.39 | 1.94 | 2.14 | 65.96 | 3.94E+08 | 8.00 | 63.46 |
| ↑↑ | Polymeric immunoglobulin receptor | Pigr | PIGR | 0.15 | -0.87 | 1.23 | 1.48 | 1.42 | 323.31 | 9.00E+08 | 18.00 | 85.00 |
| No | Lysozyme C-1 | Lyz2 | LYZ | -1.72 | -1.89 | -0.95 | 0.07 | 0.16 | 11.14 | 1.51E+07 | 2.00 | 16.79 |
| **Pro-inflammatory cytokines – ELISA** | | | | | | | | | | | | |
| **Fold-change** | **Protein names** | **Mouse Symbol** | **Human Symbol** | **Day 2** | **Day 3** | **Day 4** | **Day 5** | **Day 8** | **Score** | **Intensity** | **Peptides** | **Mol. weight [kDa]** |
| ↑↑ | Interleukin 1 beta | Il1b | IL1B | 0.24 | 0.63 | 0.03 | 0.66 | 1.7 | N/A | N/A | N/A | N/A |
| ↑↑ | Interleukin 6 | Il6 | IL6 | 0.64 | 0.79 | 0.06 | 1.21 | 1.87 | N/A | N/A | N/A | N/A |
| ↑↑↑ | Interferon gamma | Ifng | IFNG | 0.29 | 0.38 | 0.1 | 1.28 | 2.24 | N/A | N/A | N/A | N/A |
| ↑↑↑ | Tumor necrosis factor | Tnf | TNF | -0.01 | -0.05 | 0.2 | 1.99 | 2.7 | N/A | N/A | N/A | N/A |
| **Intrinsic Coagulation Pathway** | | | | | | | | | | | | |
| **Fold-change** | **Protein names** | **Mouse Symbol** | **Human Symbol** | **Day 2** | **Day 3** | **Day 4** | **Day 5** | **Day 8** | **Score** | **Intensity** | **Peptides** | **Mol. weight [kDa]** |
| No | Coagulation factor X | F10 | F10 | -0.09 | -0.43 | -0.10 | -0.39 | -0.40 | 323.31 | 5.79E+09 | 15.00 | 54.02 |
| ↓ | Coagulation factor XIII A chain | F13a1 | F13A1 | -0.01 | -0.43 | -0.54 | -1.00 | -0.68 | 118.25 | 5.34E+08 | 8.00 | 83.21 |
| ↓↓ | Coagulation factor XIII B chain | F13b | F13B | 0.01 | -0.46 | -1.00 | -1.61 | -1.54 | 323.31 | 1.06E+09 | 20.00 | 76.20 |
| No | Prothrombin | F2 | F2 | -0.06 | -0.31 | -0.17 | -0.67 | -0.41 | 323.31 | 4.68E+10 | 31.00 | 70.27 |
| ↑ | Coagulation factor V | F5 | F5 | -0.12 | -0.62 | 0.83 | 0.61 | 0.59 | 323.31 | 3.13E+09 | 32.00 | 247.23 |
| No | Coagulation factor VII | F7 | F7 | -0.21 | -0.42 | 0.57 | 0.10 | 0.26 | 61.80 | 4.11E+08 | 8.00 | 50.28 |
| ↑↑ | Fibrinogen alpha chain | Fga | FGA | 0.49 | 0.06 | 0.10 | 1.13 | 1.35 | 323.31 | 1.87E+10 | 20.00 | 87.43 |
| ↑↑↑ | Fibrinogen beta chain | Fgb | FGB | 0.55 | -0.48 | 2.35 | 2.00 | 2.07 | 288.57 | 4.39E+08 | 9.00 | 54.75 |
| ↑↑ | Fibrinogen gamma chain | Fgg | FGG | 0.25 | -0.40 | 2.08 | 1.70 | 1.85 | 58.82 | 3.75E+08 | 9.00 | 49.39 |
| ↑↑↑ | Fibrinogen-like protein 1 | Fgl1 | FGL1 | -0.13 | -0.37 | 2.49 | 1.99 | 2.21 | 45.53 | 1.29E+08 | 4 | 36.439 |
| ↓↓ | Plasminogen | Plg | PLG | -0.14 | -0.54 | -0.51 | -1.06 | -1.17 | 323.31 | 8.36E+10 | 65.00 | 90.81 |
| **Extrinsic Coagulation Pathway** | | | | | | | | | | | | |
| **Fold-change** | **Protein names** | **Mouse Symbol** | **Human Symbol** | **Day 2** | **Day 3** | **Day 4** | **Day 5** | **Day 8** | **Score** | **Intensity** | **Peptides** | **Mol. weight [kDa]** |
| ↓ | Angiotensin-converting enzyme | Ace | ACE | -0.36 | -0.88 | -0.54 | -0.77 | -0.71 | 111.36 | 4.20E+08 | 15.00 | 150.92 |
| No | Angiotensinogen | Agt | AGT | -0.06 | -0.68 | 0.04 | -0.05 | -0.10 | 323.31 | 5.23E+09 | 16.00 | 51.99 |
| ↑↑ | Coagulation factor XI | F11 | F11 | -0.13 | -0.71 | 1.90 | 1.53 | 1.93 | 38.36 | 1.65E+08 | 6.00 | 69.79 |
| ↑ | Coagulation factor XII | F12 | F12 | 0.12 | -0.27 | 1.03 | 0.12 | 0.57 | 247.11 | 4.46E+08 | 9.00 | 65.70 |
| ↑ | Coagulation factor IX | F9 | F9 | 0.02 | -0.32 | 0.70 | 0.56 | 0.62 | 248.77 | 8.84E+08 | 11.00 | 52.98 |
| ↑ | Fibronectin | Fn1 | FN1 | -0.02 | -0.27 | 1.10 | 0.91 | 0.65 | 323.31 | 1.87E+11 | 96.00 | 272.53 |
| ↓↓ | Platelet glycoprotein V | Gp5 | GP5 | 0.14 | -0.33 | -0.21 | -0.98 | -0.99 | 44.31 | 3.08E+08 | 4.00 | 63.47 |
| ↓ | Kallikrein-1 | Klk1 | KLK1 | -1.11 | -2.01 | -0.93 | 0.59 | -0.87 | 105.03 | 7.23E+07 | 5.00 | 28.77 |
| ↓↓ | Plasma kallikrein | Klkb1 | KLKB1 | 0.30 | -0.20 | -0.70 | -1.94 | -1.69 | 323.31 | 7.53E+09 | 31.00 | 71.38 |
| No | Kininogen-1 | Kng1 | KNG1 | -0.01 | -0.31 | 0.53 | 0.37 | 0.27 | 323.31 | 2.31E+11 | 29.00 | 73.10 |
| ↓↓ | von Willebrand factor | Vwf | VWF | -0.11 | -0.38 | -0.96 | -1.43 | -1.31 | 146.74 | 3.94E+08 | 12.00 | 309.27 |
| **Anticoagulants** | | | | | | | | | | | | |
| **Fold-change** | **Protein names** | **Mouse Symbol** | **Human Symbol** | **Day 2** | **Day 3** | **Day 4** | **Day 5** | **Day 8** | **Score** | **Intensity** | **Peptides** | **Mol. weight [kDa]** |
| No | Alpha-2-antiplasmin | AAP | SERPINF2 | -0.32 | -0.61 | 0.26 | -0.45 | -0.29 | 323.31 | 4.23E+10 | 22.00 | 54.97 |
| ↓↓ | Alpha-1-antitrypsin 1-1 | AAT | SERPINA1 | -0.11 | -0.45 | -0.08 | -0.49 | -0.59 | 323.31 | 2.03E+11 | 31.00 | 46.00 |
| No | Antithrombin-III | AT-III | SERPINC1 | -0.04 | -0.63 | -0.25 | -0.44 | -0.41 | 323.31 | 7.62E+10 | 41.00 | 52.00 |
| ↑↑ | Plasma protease C1 inhibitor | C1-INH | SERPING1 | -0.03 | -0.36 | 1.77 | 1.10 | 1.19 | 323.31 | 1.98E+10 | 20.00 | 55.58 |
| ↑↑ | Thrombin-activatable fibrinolysis inhibitor (TAFI) | Cpb2 | TAFI | -0.29 | -0.8 | 1.28 | 1.5 | 1.49 | 138.86 | 4.65E+09 | 9 | 48.87 |
| No | Heparin cofactor 2 | HCF2 | SERPIND1 | -0.38 | -0.83 | -0.06 | -0.51 | -0.47 | 323.31 | 6.73E+09 | 21.00 | 54.50 |
| No | Plasminogen activator inhibitor 1 | PAI-1 | SERPINE1 | -0.62 | -0.90 | 0.87 | 0.66 | 0.25 | 18.28 | 1.44E+07 | 1.00 | 45.17 |
| No | Vitamin K-dependent protein C | Proc | PROC | -0.19 | -0.44 | 0.48 | -0.02 | -0.20 | 323.31 | 7.58E+08 | 9.00 | 51.82 |

| | | | | | | | | | | | | |
|---|---|---|---|---|---|---|---|---|---|---|---|---|
| No | Vitamin K-dependent protein S | Pros1 | PROS1 | -0.12 | -0.64 | 0.67 | 0.15 | 0.07 | 163.36 | 7.10E+08 | 12.00 | 74.93 |
| ↓ | Vitamin K-dependent protein Z | Proz | PROZ | -0.06 | -0.32 | -0.23 | -0.77 | -0.64 | 323.31 | 1.58E+09 | 7.00 | 44.30 |
| ↑↑ | Protein Z-dependent protease inhibitor | PZI | SERPINA10 | -0.33 | -0.83 | 2.17 | 1.67 | 1.68 | 323.31 | 1.24E+10 | 18.00 | 51.80 |
| ↓ | Tissue factor pathway inhibitor | Tfpi | TFPI | -0.33 | -0.25 | -0.83 | -0.80 | -0.71 | 28.37 | 1.79E+07 | 1.00 | 34.99 |
| **Complement Classical Pathway** | | | | | | | | | | | | |
| **Fold-change** | **Protein names** | **Mouse Symbol** | **Human Symbol** | **Day 2** | **Day 3** | **Day 4** | **Day 5** | **Day 8** | **Score** | **Intensity** | **Peptides** | **Mol. weight [kDa]** |
| ↑↑ | Complement C1q subcomponent subunit A | C1qa | C1QA | -0.55 | -0.64 | 1.70 | 1.02 | 1.08 | 323.31 | 2.85E+09 | 7.00 | 25.97 |
| ↑↑ | Complement C1q subcomponent subunit B | C1qb | C1QB | -0.51 | -0.59 | 1.52 | 0.97 | 1.06 | 323.31 | 3.90E+09 | 5.00 | 26.72 |
| ↑ | Complement C1q subcomponent subunit C | C1qc | C1QC | -0.55 | -0.43 | 1.12 | 0.77 | 0.76 | 323.31 | 2.20E+09 | 5.00 | 25.99 |
| No | Complement C1r-A subcomponent chain | C1ra | C1R | -0.25 | -0.74 | 1.09 | 0.28 | 0.42 | 323.31 | 5.13E+09 | 23.00 | 80.07 |
| ↑ | Complement C1s-A subcomponent | C1sa | C1S | -0.09 | -0.70 | 1.36 | 0.67 | 0.84 | 323.31 | 5.35E+09 | 27.00 | 76.86 |
| No | Complement C1s-B subcomponent | C1sb | C1B | -0.21 | -0.66 | 0.25 | -0.36 | -0.14 | 26.05 | 2.09E+08 | 11.00 | 76.70 |
| **Alternative Pathway Activators** | | | | | | | | | | | | |
| **Fold-change** | **Protein names** | **Mouse Symbol** | **Human Symbol** | **Day 2** | **Day 3** | **Day 4** | **Day 5** | **Day 8** | **Score** | **Intensity** | **Peptides** | **Mol. weight [kDa]** |
| ↑↑ | Complement factor B | Cfb | CFB | -0.01 | -0.41 | 1.64 | 1.17 | 1.33 | 323.31 | 6.45E+10 | 41.00 | 85.00 |
| No | Complement factor D | Cfd | CFD | -0.25 | -0.29 | -0.04 | 0.15 | -0.10 | 323.31 | 1.92E+10 | 11.00 | 28.06 |
| **Lectin Pathway Activators** | | | | | | | | | | | | |
| Fold-change | **Protein names** | **Mouse Symbol** | **Human Symbol** | **Day 2** | **Day 3** | **Day 4** | **Day 5** | **Day 8** | **Score** | **Intensity** | **Peptides** | **Mol. weight [kDa]** |
| No | Complement component C1q receptor | Cd93 | CD93 | -0.82 | -1.35 | -0.34 | -0.05 | -0.16 | 6.56 | 6.89E+06 | 1.00 | 69.35 |
| ↓ | Ficolin-1 | Fcn1 | FCN1 | -0.07 | -0.50 | -0.50 | -0.66 | -0.72 | 251.13 | 2.26E+09 | 7.00 | 36.30 |
| ↓ | Mannan-binding lectin serine protease 1 | Masp1 | MASP1 | -0.11 | -0.49 | -0.20 | -0.39 | -0.51 | 323.31 | 1.31E+09 | 17.00 | 79.97 |
| ↓ | Mannan-binding lectin serine protease 2 | Masp2 | MASP2 | -0.17 | -0.68 | -0.23 | -0.60 | -0.52 | 229.86 | 1.27E+09 | 17.00 | 75.52 |
| ↓↓ | Mannose-binding protein C | Mbl2 | MBL2 | -0.20 | -0.35 | -0.43 | -1.56 | -1.64 | 323.31 | 5.16E+09 | 9.00 | 25.96 |
| **C3_C5-convertases** | | | | | | | | | | | | |
| **Fold-change** | **Protein names** | **Mouse Symbol** | **Human Symbol** | **Day 2** | **Day 3** | **Day 4** | **Day 5** | **Day 8** | **Score** | **Intensity** | **Peptides** | **Mol. weight [kDa]** |
| ↑↑ | Complement C2 | C2 | C2 | 0.03 | -0.74 | 1.82 | 1.35 | 1.44 | 323.31 | 4.60E+09 | 27.00 | 84.74 |
| ↑↑ | Complement C3 | C3 | C3 | -0.15 | -0.30 | 1.50 | 0.92 | 1.19 | 323.31 | 1.14E+12 | 133.00 | 186.48 |
| ↑↑ | Complement C4-B | C4b | C4B | -0.15 | -0.54 | 1.38 | 0.76 | 0.96 | 323.31 | 9.13E+10 | 86.00 | 192.91 |
| No | Complement C5 | C5 | C5 | 0.05 | -0.55 | 0.32 | -0.45 | -0.32 | 323.31 | 3.20E+10 | 72.00 | 188.88 |
| **Terminal Pathway** | | | | | | | | | | | | |
| **Fold-change** | **Protein names** | **Mouse Symbol** | **Human Symbol** | **Day 2** | **Day 3** | **Day 4** | **Day 5** | **Day 8** | **Score** | **Intensity** | **Peptides** | **Mol. weight [kDa]** |
| ↓↓ | Complement component C8 alpha chain | C8a | C8A | -0.02 | -0.78 | -0.89 | -1.64 | -1.58 | 323.31 | 1.91E+10 | 32.00 | 66.08 |
| ↓↓ | Complement component C8 beta chain | C8b | C8B | 0.02 | -0.75 | -0.59 | -1.39 | -1.28 | 323.31 | 1.15E+10 | 27.00 | 66.23 |
| ↓↓ | Complement component C8 gamma chain | C8g | C8G | -0.17 | -0.81 | -0.59 | -1.28 | -1.13 | 323.31 | 8.39E+09 | 10.00 | 22.51 |
| **No** | **Complement component C9** | **C9** | **C9** | **0.03** | **-0.84** | **-0.18** | **-0.44** | **-0.46** | **323.31** | **1.65E+10** | **32.00** | **62.00** |
| **Complement Pathway Regulators** | | | | | | | | | | | | |
| **Fold-change** | **Protein names** | **Mouse Symbol** | **Human Symbol** | **Day 2** | **Day 3** | **Day 4** | **Day 5** | **Day 8** | **Score** | **Intensity** | **Peptides** | **Mol. weight [kDa]** |
| No | C4b-binding protein | C4bpa | C4BP | 0.17 | -0.14 | 0.62 | 0.33 | 0.36 | 251.04 | 3.84E+09 | 12.00 | 51.52 |
| ↑↑ | Complement factor H | Cfh | CFH | 0.54 | -0.02 | 1.86 | 0.95 | 1.45 | 323.31 | 3.63E+10 | 51.00 | 139.14 |
| ↑ | Clusterin | Clu | CLU | -0.13 | -0.50 | 1.25 | 1.07 | 0.86 | 323.31 | 7.75E+10 | 23.00 | 51.66 |
| ↓ | Complement factor I | Cfi | CFI | -0.01 | -0.43 | -0.38 | -0.82 | -0.79 | 323.31 | 1.83E+10 | 29.00 | 67.26 |
| ↓ | Properdin | Cfp | CFP | 0.01 | -0.34 | -0.26 | -0.57 | -0.62 | 116.21 | 1.94E+09 | 8.00 | 50.33 |
| ↓ | Vitronectin | Vtn | VTN | -0.36 | -0.29 | -0.24 | -0.58 | -0.88 | 323.31 | 1.78E+10 | 11.00 | 54.85 |
| **Adverse outcome indicators in human sepsis** | | | | | | | | | | | | |
| **Fold-change** | **Protein names** | **Mouse Symbol** | **Human Symbol** | **Day 2** | **Day 3** | **Day 4** | **Day 5** | **Day 8** | **Score** | **Intensity** | **Peptides** | **Mol. weight [kDa]** |
| No | Adiponectin | Adipoq | ADIPOQ | -0.31 | -0.30 | -0.03 | -0.04 | 0.12 | 306.77 | 5.45E+09 | 5.00 | 26.81 |
| ↓↓↓ | Retinol-binding protein 4 | Rbp4 | RBP4 | 0.06 | -0.41 | -2.43 | -2.58 | -2.63 | 323.31 | 5.77E+09 | 11.00 | 23.21 |
| ↓↓ | Apolipoprotein A-I | Apoa1 | APOA1 | -0.13 | -0.52 | -0.40 | -1.32 | -1.35 | 323.31 | 1.38E+12 | 30.00 | 30.62 |
| ↓↓ | Apolipoprotein M | Apom | APOM | 0.35 | -0.44 | -0.43 | -1.08 | -1.16 | 181.52 | 3.11E+09 | 7.00 | 21.27 |
| ↓ | Gelsolin | Gsn | GSN | -0.09 | -0.47 | -0.21 | -0.87 | -0.63 | 323.31 | 6.05E+10 | 42.00 | 85.94 |
| ↓↓ | Mannose-binding protein C | Mbl2 | MBL2 | -0.20 | -0.35 | -0.43 | -1.56 | -1.64 | 323.31 | 5.16E+09 | 9.00 | 25.96 |
| ↓↓ | Corticosteroid Binding Globulin | Cbg | SERPINA6 | -0.08 | -0.34 | -0.45 | -1.56 | -1.31 | 303.41 | 9.96E+09 | 14 | 44.769 |
| **Platelet Activation** | | | | | | | | | | | | |

| Fold-change | Protein names | Mouse Symbol | Human Symbol | Day 2 | Day 3 | Day 4 | Day 5 | Day 8 | Score | Intensity | Peptides | Mol. weight [kDa] |
|---|---|---|---|---|---|---|---|---|---|---|---|---|
| ↑↑↑ | Platelet-activating factor acetylhydrolase | Pla2g7 | PLA2G7 | -0.14 | -0.64 | 2.34 | 2.91 | 2.81 | 323.31 | 5.29E+09 | 19.00 | 49.26 |

**PPSS proteins not found**

| Fold-change | Protein Name | References | Mouse Symbol | Human Symbol | Day 2 | Day 3 | Day 4 | Day 5 | Day 8 | Score | Intensity | Peptides | Mol. weight [kDa] |
|---|---|---|---|---|---|---|---|---|---|---|---|---|---|
| No | Angiopoietin-related protein 3 | | Angptl3 | ANGPTL3 | n.d. | n.d. | n.d. | n.d. | -0.24 | 13.844 | 1.71E+08 | 4 | 52.542 |
| ↑↑ | C-C motif chemokine 8 | | Ccl8 | CCL8 | n.d. | n.d. | n.d. | n.d. | 1.96 | 63.695 | 3.14E+08 | 5 | 11.017 |
| ↑ | Macrophage colony-stimulating factor 1 | | Csf1 | CSF1 | n.d. | n.d. | n.d. | n.d. | 0.65 | 5.3901 | 7.03E+07 | 4 | 60.648 |
| ↓↓ | Insulin-like growth factor I | | Igf1 | IGF1 | n.d. | n.d. | n.d. | n.d. | -1.59 | 27.959 | 7.66E+08 | 2 | 17.093 |
| ↑↑↑ | Insulin-like growth factor-binding protein 1 | | Igfbp1 | IGFB1 | n.d. | n.d. | n.d. | n.d. | 4.27 | 81.638 | 1.33E+09 | 11 | 29.57 |
| No | Insulin-like growth factor-binding protein 2 | | Igfbp2 | IGFB2 | n.d. | n.d. | n.d. | n.d. | 0.30 | 162.93 | 2.82E+06 | 9 | 32.846 |
| ↓↓ | Insulin-like growth factor-binding protein 3 | | Igfbp3 | IGFB3 | n.d. | n.d. | n.d. | n.d. | -1.73 | 37.011 | 1.38E+09 | 12 | 31.687 |
| No | Insulin-like growth factor-binding protein 4 | | Igfbp4 | IGFB4 | n.d. | n.d. | n.d. | n.d. | 0.07 | 48.659 | 3.94E+08 | 5 | 27.807 |
| No | Insulin-like growth factor-binding protein 5 | | Igfbp5 | IGFB5 | n.d. | n.d. | n.d. | n.d. | -0.12 | 34.752 | 3.45E+07 | 5 | 30.372 |
| ↓↓ | Platelet factor 4 | | Pf4 | PF4 | n.d. | n.d. | n.d. | n.d. | -1.60 | 9.5227 | 1.56E+08 | 3 | 11.243 |
| ↑↑ | Peptidoglycan recognition protein 1 | | Pglyrp1 | PGLYRP1 | n.d. | n.d. | n.d. | n.d. | 1.23 | 11.85 | 4.21E+07 | 4 | 20.489 |
| No | E-selectin | | Sele | SELE | n.d. | n.d. | n.d. | n.d. | 0.22 | 10.096 | 2.31E+08 | 4 | 66.749 |
| ↑↑ | P-selectin | | Selp | SELP | n.d. | n.d. | n.d. | n.d. | 1.00 | 51.671 | 6.52E+07 | 3 | 66.749 |
| ↑↑ | Tyrosine-protein kinase receptor Tie-1 | | Tie1 | TIE1 | n.d. | n.d. | n.d. | n.d. | 1.17 | 3.2458 | 1.27E+07 | 1 | 124.58 |
